# TumFlow: An AI Model for Predicting New Anticancer Molecules

**DOI:** 10.1101/2024.02.06.579053

**Authors:** Davide Rigoni, Sachithra Yaddehige, Nicoletta Bianchi, Alessandro Sperduti, Stefano Moro, Cristian Taccioli

**Affiliations:** Molecular Modelling Section (MMS), Department of Pharmaceutical and Pharmacological Sciences, University of Padova, Via Marzolo 5, 35131, Padova, Italy; Department of Animal Medicine, Production and Health, University of Padova, Viale dell’Università 16, 35020, Legnaro, Italy; Department of Translational Medicine, University of Ferrara, Via Luigi Borsari 46, 44121, Ferrara, Italy; Department of Mathematics “Tullio Levi-Civita”, University of Padova, Via Trieste 63, 35131, Padova, Italy

**Keywords:** Generative Model, Anticancer Molecules, Melanoma, SK-MEL-28

## Abstract

**Motivation:** Melanoma is a severe form of skin cancer increasing globally with about 324.000 cases in 2020, making it the fifth most common cancer in the United States. Conventional drug discovery methods face limitations due to the inherently time consuming and costly. However, the emergence of artificial intelligence (AI) has opened up new possibilities. AI models can effectively simulate and evaluate the properties of a vast number of potential drug candidates, substantially reducing the time and resources required by traditional drug discovery processes. In this context, the development of AI normalizing flow models, employing machine learning techniques to create new molecular structures, holds great promise for accelerating the discovery of effective anticancer therapies.

**Results:** This manuscript introduces a novel AI model, named *TumFlow*, aimed at generating new molecular entities with potential therapeutic value in cancer treatment. It has been trained on the comprehensive NCI-60 dataset, encompassing thousands of molecules tested across 60 tumour cell lines, with a specific emphasis on the melanoma SK-MEL-28 cell line. The model successfully generated new molecules with predicted improved efficacy in inhibiting tumour growth while being synthetically feasible. This represents a significant advancement over conventional generative models, which often produce molecules that are challenging or impossible to synthesize. Furthermore, *TumFlow* has also been utilized to optimize molecules known for their efficacy in clinical melanoma treatments. This led to the creation of novel molecules with a predicted enhanced likelihood of effectiveness against melanoma, currently undocumented on PubChem.

**Availability and Implementation:** https://github.com/drigoni/TumFlow.

**Supplementary information:** Uploaded.

## 1. Introduction

Melanoma, a serious form of skin cancer, originates from melanocytes which are cells responsible for producing melanin that colours the skin. It stands as the most severe type of skin cancer due to its potential to metastasize to other body parts if not detected and treated promptly. Individuals with fair skin, blue eyes, and light-coloured hair are predominantly at higher risk, largely due to their lower levels of melanin, making their skin more susceptible to harmful ultraviolet (UV) radiation from the sun (Dzwierzynski, 2021; O’Neill and Scoggins, 2019; Gandini et al., 2005; Arnold et al., 2018; Erdei and Torres, 2010). Moreover, melanoma poses an increased threat due to its resistance to conventional chemotherapy (Arioka et al., 2017). Current treatment strategies for melanoma include surgical excision, targeted therapy, and immunotherapy. Targeted therapies are employed for melanomas with specific genetic mutations, such as the BRAF V600E mutation, using inhibitors like vemurafenib and dabrafenib (Chapman et al., 2011). Immunotherapy, leveraging agents such as anti-PD-1 antibodies (nivolumab and pembrolizumab) and anti-CTLA-4 antibodies (ipilimumab), has shown efficacy in enhancing the immune response against melanoma cancer cells (Leach et al., 1996; Hodi et al., 2010; Robert et al., 2015). The drug discovery and design process are complex and resource intensive, often extending over 10-20 years with costs exceeding $2 billion (Zang and Wang, 2020; Harrer et al., 2019). Figure S1 presents the number of FDA approved drugs per year, highlighting the small increase in approvals, despite every year increasing investments in research and development visible in Figure S2.

In this context, Artificial Intelligence (AI) provides a promising avenue for revolutionizing the field, potentially reducing costs and increasing efficiency. It has become a pivotal tool in various aspects of cancer management, encompassing early detection, precision medicine, imaging, and drug repurposing (Jiang et al., 2017). Despite these advancements, the complete potential of AI in synthesizing novel anticancer molecules is yet to be fully harnessed and explored (Vamathevan et al., 2019). Within AI, Machine Learning and Deep Learning are key subfields, including techniques like supervised and unsupervised learning. Supervised learning is utilized for tasks like disease detection and drug efficiency estimation, while unsupervised learning aids in patient stratification and disease recognition (Hassanzadeh et al., 2019). Deep Learning, particularly effective in processing large datasets such as the use of images, has contributed notably to melanoma cancer diagnostics among other areas (Munir et al., 2019).

Among the unsupervised models, there is a family of approaches that fall under the name of generative models, which are a class of algorithms designed to learn and generate new data that is similar to a given training dataset. These models aim to capture the underlying patterns and structures in the training data, enabling them to generate novel samples that share characteristics with the original data. In the field of new drug generations, various approaches based on Variational Autoencoders (VAEs) (Kingma and Welling, 2013; Rigoni et al., 2020a,b, 2023; Hy and Kondor, 2023), Generative Adversarial Networks (GANs) (De Cao and Kipf, 2018; Tsujimoto et al., 2021), Normalizing Flows (Shi et al., 2019; Zang and Wang, 2020; Faez et al., 2021) and Diffusion Models (Vignac et al., 2022; Xu et al., 2022; Huang et al., 2023) have been explored. Moreover, the advent of large language models (LLMs) using transformer architectures (Vaswani et al., 2017) has further expanded this field. Transformer-based models, originally developed for natural language processing tasks, have been successful in capturing complex patterns in data and have been applied in drug generation (Mazuz et al., 2023; Bagal et al., 2021). Generative models can also include a predictive model to predict the antitumoral activity of generated molecules, enabling the identification of the most promising candidates.

Normalizing flow methods have been applied in various fields, including density estimation (Huang et al., 2018) and data augmentation. They have been used in tasks such as image generation, speech synthesis, and molecular generation in chemistry. Specific architectures like Real Non-Volume Preserving (RealNVP) (Dinh et al., 2016) and Glow (Kingma and Dhariwal, 2018) are examples of these methods. The normalizing flow method represents an effective technique to learn the unknown probability distribution that has generated the data in the training set, i.e. the chemical structure of the molecules. It does it by employing a series of invertible transformations to transmute a probability distribution over input data (i.e., molecule structures) into a designated target probability distribution.

This research incorporates Deep Learning into drug discovery with *TumFlow*, a novel approach for generating molecular graphs for cancer therapeutics. *TumFlow*, Building on the foundational work of MoFlow (Zang and Wang, 2020), a pioneering model in the field of models applied to graph structures, adapts and enhances these capabilities specifically to address melanoma treatment challenges. It leverages MoFlow efficient bond and atom generation to create novel molecules aimed to be effective against melanoma cancer cells. When learning to generate new antitumoral molecules, *TumFlow* is trying to solve a complex assignment made of challenging subtasks. The successful generation of useful molecules requires an implicit comprehension of their pharmacokinetics, the identification of single or multiple targets, and the assurance that they bind with high affinity to these to inhibit tumour progression. Each of these subtasks presents formidable difficulties independently, and the fact that the neural network does not have this kind of information to learn from makes the learning process even more challenging.

The integration of *TumFlow* into the drug discovery process reflects a broader trend in AI increasing impact on healthcare and pharmaceutical research. By focusing specifically on melanoma, *TumFlow* addresses a critical need in cancer treatment, offering the potential to rapidly identify and develop new therapeutic molecules. For this reason, this work represents a novel contribution to anticancer drug discovery.

## 2. Methods

*TumFlow* is based on MoFlow, a normalizing flow model originally developed for the generation of graph structures without any focus in anticancer molecules. On the other hands, *TumFlow* aims to learn the unknown probability distribution that has generated the chemical structures of the molecules in the dataset, with the purpose of using the learned distribution to generate new novel chemical structures that should convey similar substructures and similar anticancer activities. Therefore, *TumFlow* is developed to predict new antitumor molecules against the SK-MEL-28 melanoma, addressing all the unique challenges and requirements of melanoma treatment. It is trained on the comprehensive NCI-60 dataset, made public by the National Cancer Institute^1^, which encompasses thousands of molecules tested across a broad spectrum of tumour cell lines.

The following sections will present more in detail the data preprocessing method applied to the NCI-60 dataset, as well as, the *TumFlow* model. Additional details about Normalizing Flows are reported in Section S3 of the Supplementary Material, while more details on the *TumFlow* model are reported in Section S4.

The following mathematical notation is adopted: (i) lower case symbols for scalars, indexes, and assignment to random variables, e.g., *n* and *x*; (ii) italics upper case symbols for sets and single random variables, e.g., *A* and *X*; (iii) bold lower case symbols for vectors and assignments to vectors of random variables, e.g., ***a*** and ***x***; (iv) bold upper case symbols for matrices, tensors, and vectors of random variables, e.g., ***A*** and ***Z***; (v) the position within a tensor or vector is denoted by numeric subscripts in square brackets, for example, ***A***_[*i,a*:*b*,:]_ where *i, a, b ∈ ℕ* ^+^, and “:” indicates the positions from *a* to *b*. The solitary use of the colon symbol “:” represents all positions. (vi) calligraphic symbols for domains, e.g., *Q*; (vii) when it is clear from the context, the probability random variables are omitted, as 𝕡 (*x*) instead of 𝕡 (*X* = *x*).

### 2.1. Data Sources and Data Preprocessing

The NCI-60 project^2^, launched in 1990, employs 60 human tumor cell lines representing diverse cancers to evaluate up to 7,000 small molecules annually for anticancer properties.

It provides four files with the results of their experiments: “GI50.csv”, “LC50.csv”, “IC50.csv”, and “TGI.csv”. In this work, only the “GI50.csv” file was used. It contains data on the GI50 (Growth Inhibition of 50%) values, which are derived from laboratory assays that measured the concentration of a chemical compound required to inhibit the growth of a specific tumor cell line by 50%. These values serve as a direct indicator of the compound potential antitumoral efficacy. The dataset is structured with various fields, including the National Service Center (NSC) code, a unique numeric identifier assigned to substances tested and evaluated by the National Cancer Institute. It also contains information on the tested cell line and the respective GI50^3^ value. The data was acquired through standardized procedures to ensure the consistency and reliability of the measurements. A lower GI50 value indicates a higher efficacy of the molecule in inhibiting tumour growth in the tested cells.

The choice of this file was based on its relevance in identifying compounds with potential antitumoral efficacy, particularly in the context of SK-MEL-28 melanoma cells. In fact, a preliminary data analysis visible in Figure S3 revealed that molecules used clinically *show better representation* in the GI50 dataset. Indeed, the GI50 scores offer a more accurate representation of clinical drugs since they exhibit a more evenly distributed pattern and better distinguish the effects of various drugs. This correlation reinforces the validity of the approach presented in this work and highlights the importance of integrating real and clinically relevant data into the modelling process.

The training of the *TumFlow* model was performed on this data, focusing only on molecules tested on SK-MEL-28 melanoma cell lines made by chemical elements commonly found in organic compounds^4^. During the data preprocessing phase, all molecules with a positively charged oxygen (O^+^) were removed, and all “ion pair” compounds were sanitized selecting only the largest connected component as the main molecule structure while discarding the remaining smaller component(s). If the sanitization resulted in a structure already existing in the dataset, only the experiment with the highest efficacy score was retained. For molecules with multiple in-vitro experiments, the corresponding GI50 values were averaged. Following the data preprocessing phase, the dataset consists of 46.766 unique molecules, each paired with its corresponding GI50 efficacy value.

### 2.2. TumFlow

*TumFlow* aims to predict new anticancer molecules exploiting the graph representation of the molecule structure, differently from other works adopting the SMILES (Weininger, 1990) sequential representation.

Mathematically, let *D* be the dataset, *T r* be the training set of molecules, Θ_*n*_ and Θ_*e*_ be respectively the set of the atom types and the set of the edge types extracted from the dataset *D*. Let *d*_*v*_ =| Θ_*n*_ | be the number of atom types, *d*_*e*_ =| Θ_*e*_ | be the number of edge types, and *d*_*n*_ the maximum number of atoms, hydrogens excluded, forming the molecules in the dataset *D*. Then, a molecule is represented as a graph *G ∈ 𝒢*:

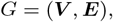

where 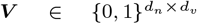 is a node type matrix and 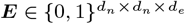 is an edge type tensor, such that ***V***_*i,v*_ = 1 only if the molecule node *i* is of type *v* and that ***E***_*i,j,e*_ = 1 only if the molecule nodes *i* and *j* are connected through a bond of type *e*. The set of all the possible graphs is defined as:

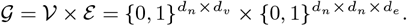

*TumFlow* aims to learn the complex probability distribution ℙ_*𝒢*_, from which the molecules in the dataset are generated, in order to sample from it new useful molecule graphs 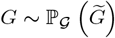.

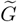 denotes the random variable over graph structures with support in *G. TumFlow* factorizes the probability distribution as:

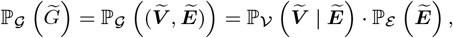

where 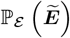 is the probability distribution over molecule bounds, 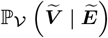 is the conditioned probability distribution over atoms given molecule bounds, and both 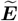 and 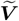 are vectors of random variables. In simpler terms, *TumFlow* first predicts the set of bonds forming the structure of the molecule and then conditions the generation of the molecule atoms by the predicted bonds. It uses two normalizing flow models jointly trained. The first is used for predicting the molecule bonds and the second is used to predict the molecule atoms. Figure 1 reports the overall model architecture summarizing the main steps.

**Fig. 1:**
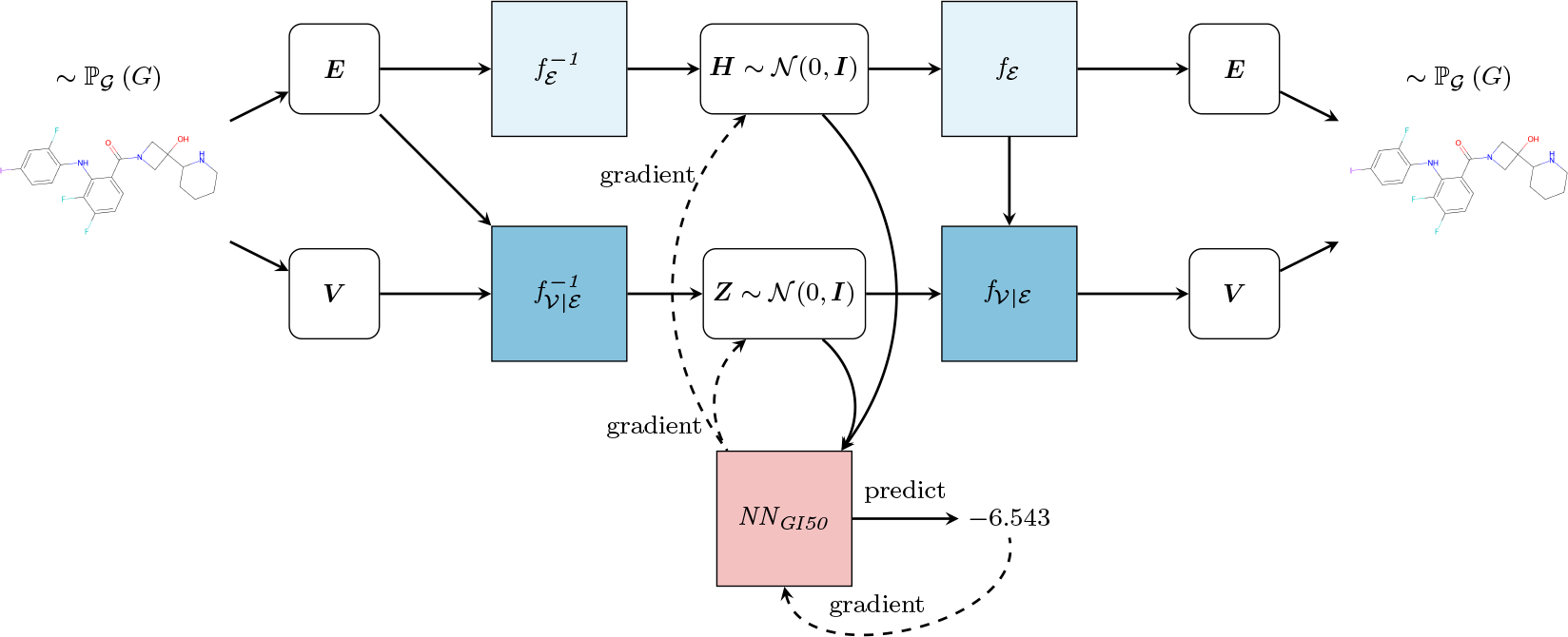
Overview of *TumFlow* model. From the molecule in input (on the left) the graph *G* = (***V***, ***E***) is constructed, and both the latent representations 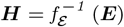 (***E***) and 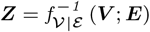 are obtained. From the latent representations, the graph *G* and then the molecule in output (on the right) are reconstructed through the functions *f*_*ε*_ and *f*_*V*|*ε*_. The module *NN*_*GI50*_ predicts the GI50 score and can optimise the molecule structure employing the gradient descend approach.

**Fig. 2:**
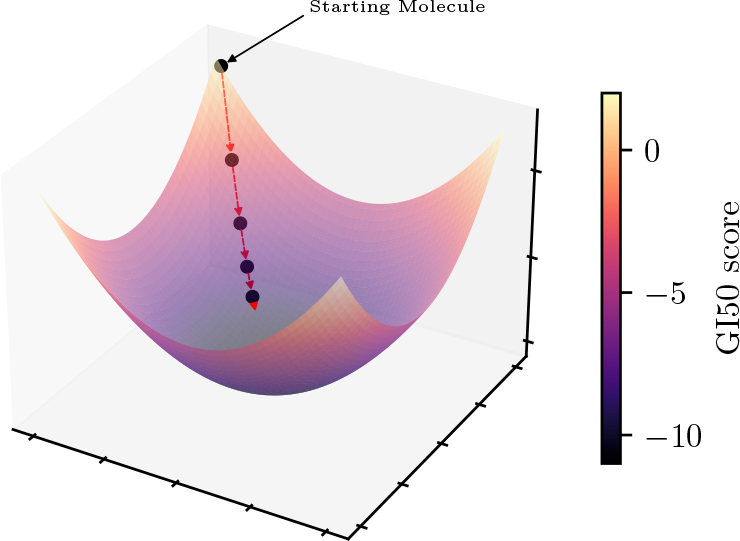
Figure representing the main idea behind the *TumFlow* generation of novel molecules with lower GI50 values, i.e., high antitumoral efficacy. Starting from an initial good molecule structure, new molecules are sampled in compliance with the gradient descent approach.

*TumFlow* is trained to optimize the negative log-likelihood loss:

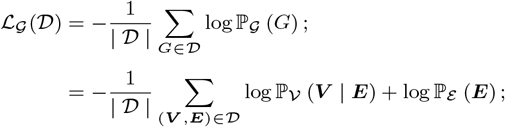

with:

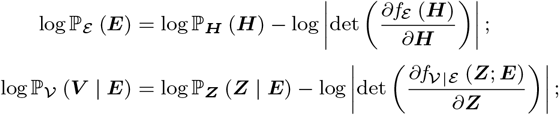

where *f*_*E*_ and *f*_*V*|*ε*_ are two invertible and differentiable functions to learn, ***H*** and ***Z*** are respectively two latent representations for atom and adjacency tensors, 𝕡 _***H***_ and 𝕡 _***Z***_ are the two simple target distributions, i.e., two standard normal distributions *𝒩* (0, ***I***), with zero mean and identity matrix as covariance matrix.

Affine coupling layers are used in the implementation of both 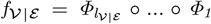 and 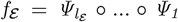, where *l*_*V*|*ε*_ and *l*_*ε*_ represents the number of coupling layers composing *f*_*V*|*ε*_ and *f*_*ε*_, respectively. For the sake of clarity, the explicit dependency on ***E*** in the notation of *Φ*_*i*_ is omitted. In the implementation of the coupling layers, the sigmoid function replaces the exponential function, as it provides better numerical stability when stacking multiple coupling layers. Mathematically, each function 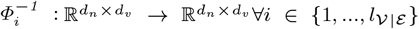 splits the input into two parts according to the node types dimension *d*_*v*_. Given 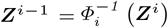 and a selected dimension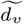:

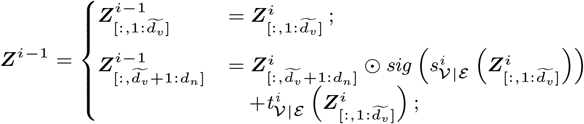

where ***Z*** = ***Z***^0^ and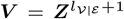. Functions 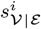 and 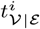 are multi-layer perceptions (MLPs) based on the output of a graph neural network (Schlichtkrull et al., 2018), which aims to learn the representation of the graph underlying the molecular structure.

Similarly, each function 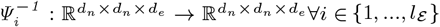 splits the input into two parts according to the bond types dimension *d*_*e*_. Given 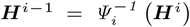 and a selected dimension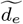:

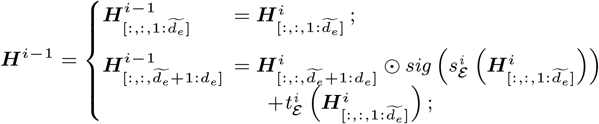

where ***H*** = ***H***^0^ and ***E*** = ***H***^*l*+1^. Functions 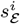 and 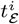 are implemented with a sequence of 2D convolutional neural network.

It is essential to consider that the normalizing flow framework is designed for continuous space values, and as such, it cannot be directly applied to discrete structures like both node and adjacency tensors. To address this limitation, a pre-processing step is implemented before the utilization of coupling layers. Specifically, a random uniform noise drawn from a carefully selected interval of values is added to each entry of the tensors. This introduction of noise serves the purpose of incorporating a continuous element into the discrete structures, aligning them with the framework requirements and enabling the subsequent application of coupling layers. The carefully selected noise distribution enables the accurate selection of corresponding atoms and bonds, through the argmax function, when utilizing *f*_*V*|*ε*_ and *f*_*ε*_ to reconstruct *G*.

#### 2.2.1. Prediction of the GI50 Scores

*TumFlow* includes a nonlinear neural network *NN*_*GI*50_ designed to predict the antitumor activity of individual molecules. This neural network is learned once the main normalizing flow networks have been learned. More precisely, the neural network is trained to predict the GI50 score associated with the molecule latent representation. Mathematically, *NN*_*GI*50_ represents the function:

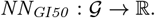

This function is trained with the mean squared error (squared L2 norm) loss computed among predicted values and in-vitro measured values. More in detail, given a molecule graph *G*, its predicted antitumor activity *p* is estimated as:

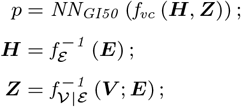

where the *f*_*vc*_ is a function that linearizes both the tensors ***H*** and ***Z***, and then concatenates them together.

#### 2.2.2. New Molecule Generation

After the learning phase, *TumFlow* can generate novel chemical structures conveying similar antitumoral activities to those in the training set and, using *NN*_*GI*50_, it can also predict the antitumoral activity for each molecule. While this approach “used alone” proves highly valuable in other research domains, such as computer vision where realistic face images need to be generated (Wu et al., 2021), it shows limitations when creating new antitumoral molecules. In fact, unlike scenarios where realistic faces are generated from datasets comprised of numerous real-world face images, the NCI-60 dataset includes many molecules with suboptimal antitumor efficacy. Consequently, employing this simplistic method to develop new molecules may yield structures with limited antitumor activities. *TumFlow* generates new molecules adopting a different approach that closely resembles structure optimization. Starting from a molecule with high antitumoral efficacy, through the use of the *NN*_*GI*50_ neural network, it modifies its structure to exhibit better antitumoral effects. Given a molecule in input, the optimization takes place many times, and for each of them, *TumFlow* predicts the new molecule structure with associated predicted GI50 scores.

The optimization, also visible in Figure S4, is performed using the gradient descent method, which is a pervasive optimization algorithm in machine learning, employed to reduce the value of an objective function. Its primary objective is to iteratively approach the minimum of a function by moving in the direction of the most significant decrease in that function. In other words, *TumFlow* applies the gradient descent technique to the function *NN*_*GI*50_ aiming to minimize the GI50 score. The optimization starts from a given molecule i.e., the black ball in the top part of the image. Then, the *NN*_*GI*50_ predicts the GI50 score associated with the starting molecules, as well as, the direction to follow in the latent space to minimize the score. In other words, the direction to follow is the negative of the gradient returned by the *NN*_*GI*50_ component as the lower the value is, the better the antitumoral properties are. Consequently, a new point in the latent space is selected, which can be decoded back to a molecule structure through *f*_*E*_ and *f*_*V*|*E*_. This process is repeated several times. *TumFlow* performs the optimization process outlined above for each molecule, taking into account various gradient descent step values. In other words, for each optimization process, the movement performed in latent space involves making jumps of varying distances.

The generation of new molecules is performed following two different approaches: *i)* in the first approach, the starting point consists of molecules with higher antitumoral efficacy appearing in the training set, i.e., antitumoral molecules tested in vitro from the NCI-60 project; while *ii)* in the second approach, the starting point consists of nine molecules, reported in Table S1, known for their efficacy in clinical treatments for melanoma.

In the process of generating new molecules, *TumFlow* has been equipped with the ability to assess the Synthetic Accessibility Score (SAS) of drug-like molecules based on molecular complexity and fragment contributions (Ertl and Schuffenhauer, 2009). This incorporation is essential because, even though *TumFlow* allows to generate novel molecules not seen before, sometimes it produces energetically unstable structures and complex molecules that are challenging to synthesize. Further details will be discussed in Section S7 of the Supplementary Material. In this work, the reported SAS values are normalized to fall within the range of [0, 1], with lower values indicating greater ease of molecule synthesis and higher values suggesting increased difficulty in the synthesis process. The incorporation of this metric significantly enhanced the quality of molecules generated by *TumFlow*, facilitating the identification of compounds with potential antitumor effectiveness as well as a desired level of synthesis complexity.

## 3. Results and Discussion

This section presents some novel molecules generated by *TumFlow* against the SK-MEL-28 melanoma tumour, while Section S7 of the Supplementary Material presents its limitations. Two processes were adopted for generating new molecules with *TumFlow* that will be individually discussed in the following. Each molecule will be introduced with its predicted GI50 score, the normalized SAS value, and its similarity measure to the initial molecule. The similarity score allows to evaluate how much the similarity between the newly generated molecule and the starting molecule differ from each other. This score is calculated using the Tanimoto similarity of the Morgan Fingerprint (Rogers and Hahn, 2010).

### 3.1. Generation Starting from the NCI-60 Dataset

This section presents a chosen set of novel molecules generated by *TumFlow* considering molecules from the training set as starting points. Specifically, in this generation procedure, the first 140 molecules with higher antitumoral efficacy appearing in the training set, i.e., antitumoral molecules tested in vitro from the NCI-60 project, were used in turn as a starting point.

Figure 3 presents some molecules obtained from a provided starting molecule, while the corresponding canonical SMILES are reported in Table S2. Specifically, the figure presents a grid where the first column on the left depicts the starting molecule structures, while the other columns report the new molecules obtained from them. By inspecting the molecular structures and the corresponding predicted GI50 scores, it is visible that *TumFlow* attempts to enhance the structure of the provided starting molecule to improve its efficacy against melanoma tumours. Nevertheless, in the process, the model tends to generate (i) increasingly complex molecules, introducing synthesis issues, and (ii) molecular structures dissimilar from the starting one.

**Fig. 3:**
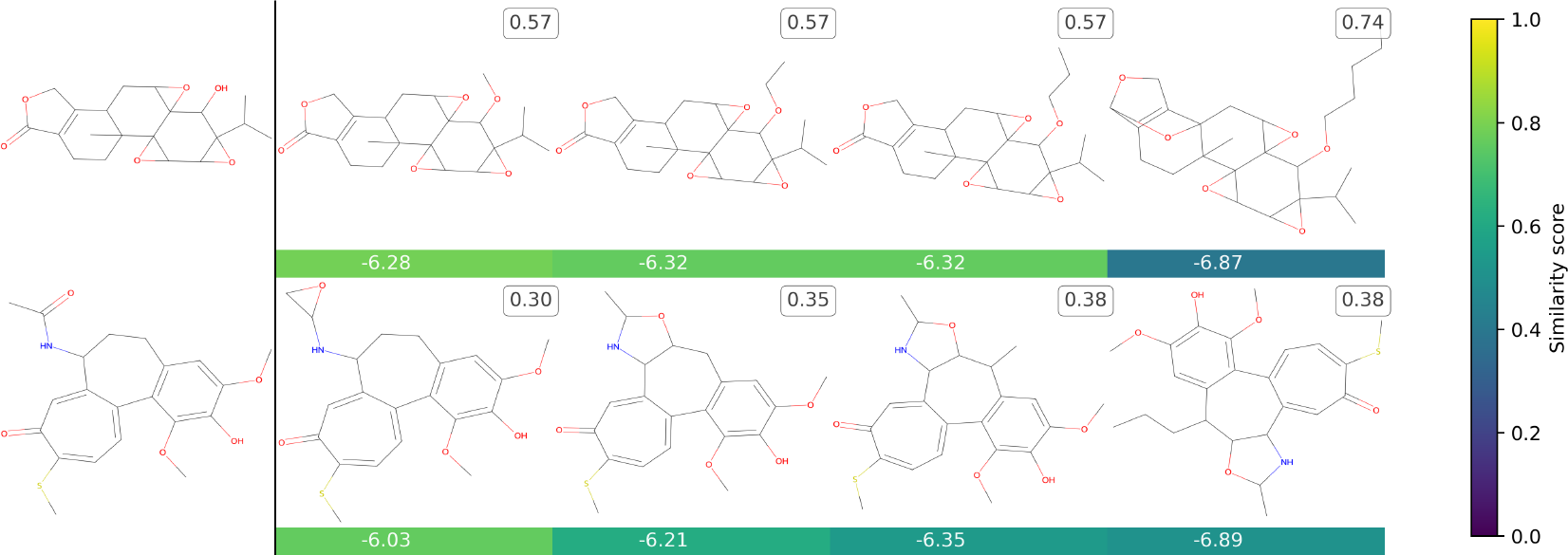
This grid presents the novel molecules that *TumFlow* has generated starting from those in the dataset. The first column on the left reports the starting molecule structures, while the other columns report the new molecules. The score reported under each generated molecule represents the *TumFlow* predicted GI50 score, while the colour conveys the similarity score of the newly generated structure with the starting molecule structure. The score on the molecule top right box reports the normalized SAS score.

Figure 4 presents more novel molecules, while the corresponding canonical SMILES, including those of the provided starting molecules, are reported in Table S3. All of these molecules are chemically noteworthy and interesting, particularly due to their absence in the dataset. It is important to notice that, the generation process sometimes results in uncommon molecules that encounter challenges in synthesis and/or contain rare substructures. Nevertheless, thanks to the SAS score, it becomes possible to identify molecules that are challenging to synthesize and, consequently, filter them as necessary.

**Fig. 4:**
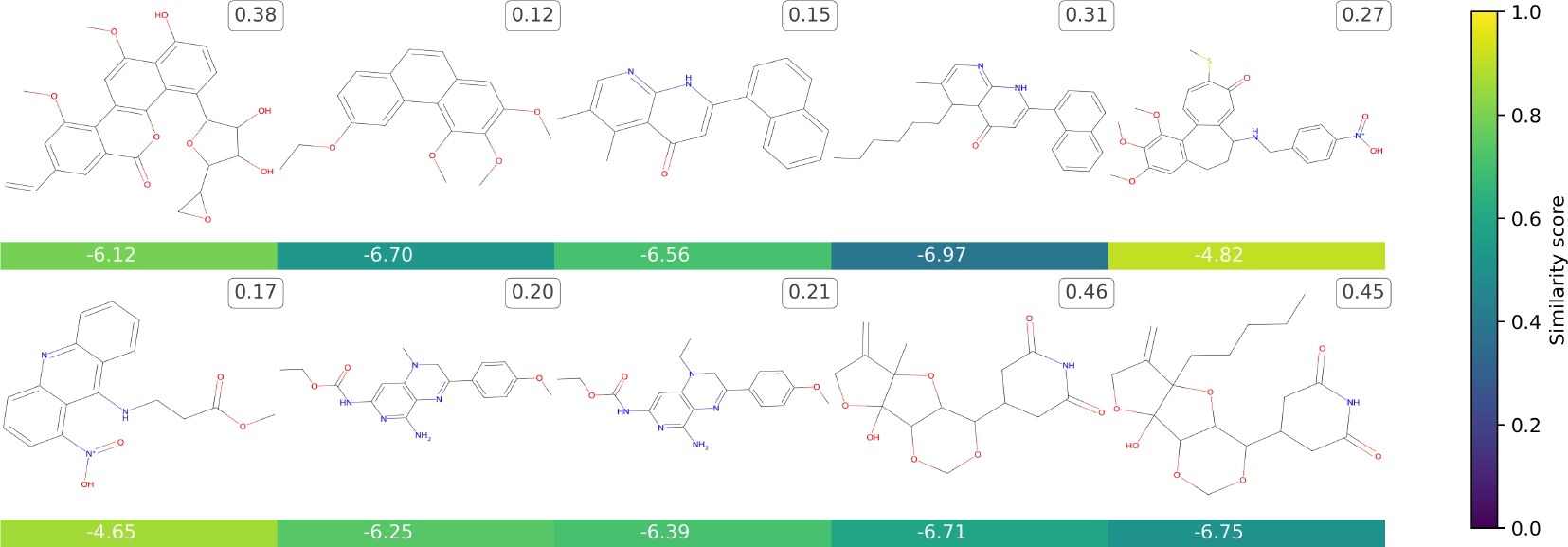
This grid presents the novel molecules that *TumFlow* has generated starting from those in the dataset. The score reported under each generated molecule represents the *TumFlow* predicted GI50 score, while the colour conveys the similarity score of the newly generated structure with the starting molecule structure. The score on the molecule top right box reports the normalized SAS score.

All the novel molecules, except the first compound^5^ generated by the first molecule reported in Figure 3, are not available on PubChem^6^. This absence of novel molecules on PubChem highlights the pioneering nature of *TumFlow* in exploring unknown chemical spaces, while the already existing molecule, demonstrates the ability of the model to generate meaningful structures.

### 3.2. Generation Starting from Clinically Adopted Anti-Melanoma Molecules

This section presents a selected set of novel molecules generated from *TumFlow* considering clinical molecules as starting points. Specifically, some of the nine molecules reported in Table S1, known for their efficacy in clinical treatments for melanoma, were used as starting points.

Figure 5 presents some novel molecules, while the corresponding canonical SMILES are reported in Table S4. Regarding the molecules generated from clinical drugs, a pattern akin to those originating from in vitro molecules is discernible. In this scenario as well, *TumFlow* demonstrates the capacity to generate novel molecular structures, albeit occasionally encountering challenges in synthesis. Notably, with the exception of just one molecule, all the newly generated structures are absent from PubChem. In fact, the initial molecule derived from the first clinical drug, specifically the second structure in the first row of the image, has been identified as a previously studied compound against cancer, bearing the corresponding “NSC = 133726”^7^. More precisely, this compound is subjected to **in vivo testing** on mice to assess its efficacy against Leukemia L1210.

**Fig. 5:**
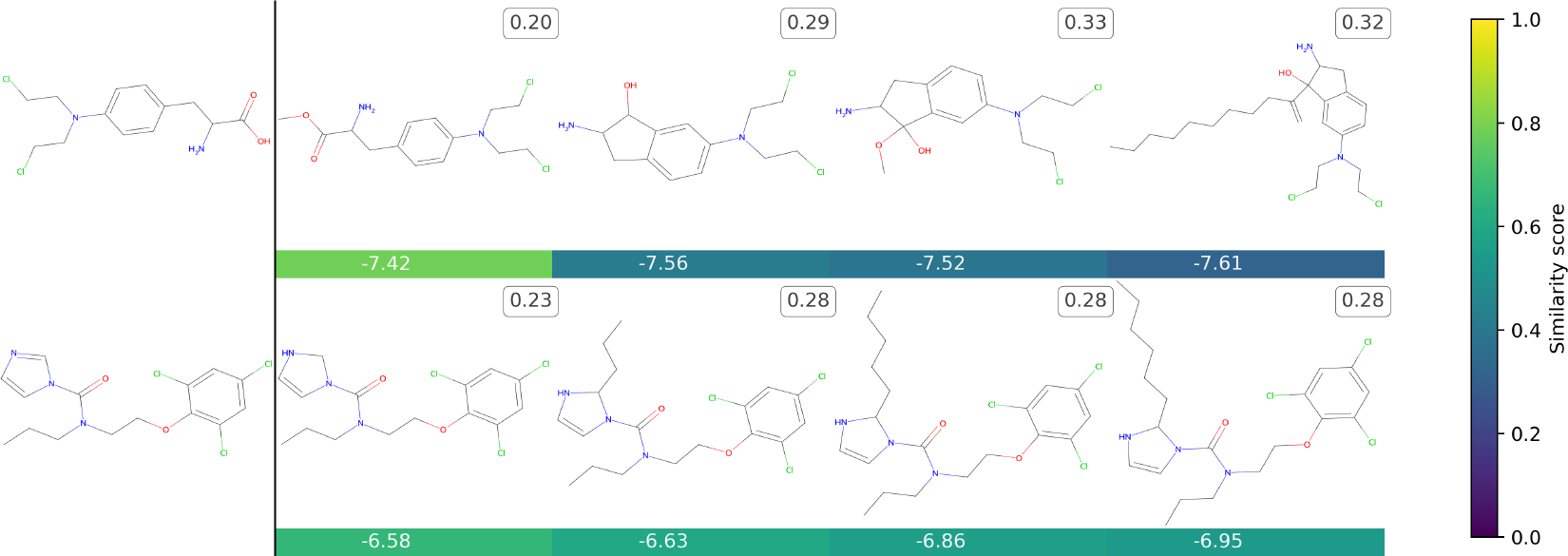
This grid presents the novel molecules that *TumFlow* has generated starting from three clinical molecules. The first column reports the starting molecule structures, while the other columns report the new molecules. The score reported under each generated molecule represents the *TumFlow* predicted GI50 score, while the colour conveys the similarity score of the newly generated structure with the starting molecule structure. The score on the molecule top-right box reports the normalized SAS score.

The identification of a novel molecule, previously studied for its anticancer properties and **not present in the training set**, underscores *TumFlow* potential to explore chemical space beyond the confines of existing datasets. This capability suggests that *TumFlow* has the capacity to propose compounds with therapeutic relevance that might not have been part of the original training data, stressing its potential to contribute to the discovery of compounds with valuable properties.

## 4. Future Prospects

Future research should consider various strategies to advance and increase the efficacy of the model presented in the context of oncological studies. One of these is the integration of broader and more diverse datasets, encompassing a wide variety of cancer types and molecules. Indeed, by exploiting drugs addressing different types of cancer, the model could learn complementary information that could enhance the discovery of new and more effective molecules. Moreover, given the *TumFlow* tendency to generate energetically unstable complex structure, future works will consider the inclusion of the SAS values during the generation process and also the inclusion of metrics to account for the energetic stability of the molecule. Additionally, incorporating inhibition, lethality and toxicity of antitumor molecules information could provide an even more accurate algorithm for predicting new anticancer drugs. Another critical aspect is the continual enhancement of the computational and algorithmic capabilities of the model, to tackle challenges like interpreting molecular mechanisms and predicting drug side effects and resistance with the purpose of optimizing the molecules to yield better properties.

## 5. Conclusion

This investigation prominently showcases the generative capabilities and potential of *TumFlow* in oncological drug development. Unlike conventional methodologies, the presented approach harnesses the distinctive strengths of normalizing flow algorithms, notably their adeptness at modelling complex molecule distributions and generating accurate new data samples. This marks a substantial leap beyond traditional AI techniques, delivering unprecedented precision and efficiency in generating novel anticancer molecules.

In particular, this work demonstrates that *TumFlow* can identify crucial patterns and correlations between molecular structures and their predicted effectiveness against tumours like melanoma. It not only exhibits creativity in generating novel and promising molecules that have not been seen before but also has the capability to generate molecules not included in the training dataset, which already exist and have been subjected to in vivo experiments for antitumoral assessment. Although this creativity is essential for generating new drugs, some limitations that come with it were presented, such as the feasibility of synthesis and the chemical instability of some generated structures.

By redefining the boundaries of possibilities within normalizing flow algorithms, the *TumFlow* model emerges not merely as a predictive instrument but as a designer, hopefully, of future oncological therapies. This methodology promises to diminish the time and financial constraints associated with drug discovery, steering researchers toward an era where swift, targeted, and potent cancer therapies are not merely conceivable but attainable. In summary, the application of the *TumFlow* model, as presented in this study, represents a significant advancement in the fight against cancer. This endeavour not only exemplifies the model’s current achievements but also paves the way for a myriad of advancements in anticancer treatments and patient care.

## Footnotes

1 https://dtp.cancer.gov/

2 https://dtp.cancer.gov/discovery_development/nci-60/

3 The GI50 values are obtained by interpolating the *GIPRCNT* scores, which are the percentage of treated cell growth as a fraction of control cell growth corrected for the count of cells at the time of drug addition in the assay. 100 is control growth, 0 is complete inhibition of growth (cytostasis), and -100 is complete cell kill. More information is reported in the NCI-60 project website.

4 Only molecules composed by hydrogen (H), carbon (C), nitrogen (N), oxygen (O), fluorine (F), phosphorus (P), sulfur (S), chlorine (Cl), selenium (Se), bromine (Br), and iodine (I).

5 CID = 121297650

6 https://pubchem.ncbi.nlm.nih.gov/

7 CID = 421441

## References

Arioka, M., Takahashi-Yanaga, F., Kubo, M., Igawa, K., Tomooka, K., and Sasaguri, T. (2017). Anti-tumor effects of differentiation-inducing factor-1 in malignant melanoma: Gsk-3-mediated inhibition of cell proliferation and gsk-3-independent suppression of cell migration and invasion. Biochemical pharmacology, 138:31–48.

Arnold, M., de Vries, E., Whiteman, D. C., Jemal, A., Bray, F., Parkin, D. M., and Soerjomataram, I. (2018). Global burden of cutaneous melanoma attributable to ultraviolet radiation in 2012. International Journal of Cancer, 143:1305–1314.

Bagal, V., Aggarwal, R., Vinod, P., and Priyakumar, U. D. (2021). Molgpt: molecular generation using a transformer-decoder model. Journal of Chemical Information and Modeling, 62(9):2064–2076.

Chapman, P. B., Hauschild, A., Robert, C., Haanen, J. B., Ascierto, P., Larkin, J., Dummer, R., Garbe, C., Testori, A., Maio, M., et al. (2011). Improved survival with vemurafenib in melanoma with braf v600e mutation. New England Journal of Medicine, 364(26):2507–2516.

De Cao, N. and Kipf, T. (2018). MolGAN: An implicit generative model for small molecular graphs. ICML 2018 workshop on Theoretical Foundations and Applications of Deep Generative Models.

Dinh, L., Sohl-Dickstein, J., and Bengio, S. (2016). Density estimation using real nvp. In International Conference on Learning Representations.

Dzwierzynski, W. W. (2021). Melanoma risk factors and prevention. Clinics in plastic surgery, 48:543–550.

Erdei, E. and Torres, S. M. (2010). A new understanding in the epidemiology of melanoma. Expert Review of Anticancer Therapy, 10:1811–1823.

Ertl, P. and Schuffenhauer, A. (2009). Estimation of synthetic accessibility score of drug-like molecules based on molecular complexity and fragment contributions. Journal of cheminformatics, 1:1–11.

Faez, F., Ommi, Y., Baghshah, M. S., and Rabiee, H. R. (2021). Deep graph generators: A survey.

Gandini, S., Sera, F., Cattaruzza, M. S., Pasquini, P., Zanetti, R., Masini, C., Boyle, P., and Melchi, C. F. (2005). Meta-analysis of risk factors for cutaneous melanoma: Iii. family history, actinic damage and phenotypic factors. European journal of cancer (Oxford, England : 1990), 41:2040–2059.

Harrer, S., Shah, P., Antony, B., and Hu, J. (2019). Artificial intelligence for clinical trial design. Trends in pharmacological sciences, 40:577–591.

Hassanzadeh, P., Atyabi, F., and Dinarvand, R. (2019). The significance of artificial intelligence in drug delivery system design. Advanced Drug Delivery Reviews.

Hodi, F. S., O’day, S. J., McDermott, D. F., Weber, R. W., Sosman, J. A., Haanen, J. B., Gonzalez, R., Robert, C., Schadendorf, D., Hassel, J. C., et al. (2010). Improved survival with ipilimumab in patients with metastatic melanoma. New England Journal of Medicine, 363(8):711–723.

Huang, C.-W., Krueger, D., Lacoste, A., and Courville, A. (2018). Neural autoregressive flows. In International Conference on Machine Learning, pages 2078–2087. PMLR.

Huang, H., Sun, L., Du, B., and Lv, W. (2023). Conditional diffusion based on discrete graph structures for molecular graph generation. volume 37.

Hy, T. S. and Kondor, R. (2023). Multiresolution equivariant graph variational autoencoder. Machine Learning: Science and Technology, 4.

Jiang, F., Jiang, Y., Zhi, H., Dong, Y., Li, H., Ma, S., Wang, Y., Dong, Q., Shen, H., and Zhao, Y. (2017). Artificial intelligence in healthcare: past, present and future. Stroke and Vascular Neurology.

Kingma, D. P. and Dhariwal, P. (2018). Glow: Generative flow with invertible 1x1 convolutions. Advances in neural information processing systems, 31.

Kingma, D. P. and Welling, M. (2013). Auto-encoding variational bayes. In Proceedings of the 30th International Conference on Machine Learning, pages 307–315.

Leach, D. R., Krummel, M. F., and Allison, J. P. (1996). Enhancement of antitumor immunity by ctla-4 blockade. Science, 271(5256):1734–1736.

Mazuz, E., Shtar, G., Shapira, B., and Rokach, L. (2023). Molecule generation using transformers and policy gradient reinforcement learning. Scientific Reports, 13(1):8799.

Munir, K., Elahi, H., Ayub, A., Frezza, F., and Rizzi, A. (2019). Cancer diagnosis using deep learning: a bibliographic review. Cancers, 11(9):1235.

O’Neill, C. H. and Scoggins, C. R. (2019). Melanoma. Journal of surgical oncology, 120:873–881.

Rigoni, D., Navarin, N., and Sperduti, A. (2020a). Conditional constrained graph variational autoencoders for molecule design.

Rigoni, D., Navarin, N., and Sperduti, A. (2020b). A systematic assessment of deep learning models for molecule generation. arXiv preprint arXiv:2008.09168.

Rigoni, D., Navarin, N., and Sperduti, A. (2023). Rgcvae: Relational graph conditioned variational autoencoder for molecule design. arXiv preprint arXiv:2305.11699.

Robert, C., Long, G. V., Brady, B., Dutriaux, C., Maio, M., Mortier, L., Hassel, J. C., Rutkowski, P., McNeil, C., Kalinka-Warzocha, E., et al. (2015). Nivolumab in previously untreated melanoma without braf mutation. New England journal of medicine, 372(4):320–330.

Rogers, D. and Hahn, M. (2010). Extended-connectivity fingerprints. Journal of chemical information and modeling, 50(5):742–754.

Schlichtkrull, M., Kipf, T. N., Bloem, P., Van Den Berg, R., Titov, I., and Welling, M. (2018). Modeling relational data with graph convolutional networks. In The Semantic Web: 15th International Conference, ESWC 2018, Heraklion, Crete, Greece, June 3–7, 2018, Proceedings 15, pages 593–607. Springer.

Shi, C., Xu, M., Zhu, Z., Zhang, W., Zhang, M., and Tang, J. (2019). Graphaf: a flow-based autoregressive model for molecular graph generation. In International Conference on Learning Representations.

Tsujimoto, Y., Hiwa, S., Nakamura, Y., Oe, Y., and Hiroyasu, T. (2021). L-molgan: an improved implicit generative model for generation of large molecular graphs. ChemRxiv.

Vamathevan, J., Clark, D., Czodrowski, P., Dunham, I., Ferran, E., Lee, G., Li, B., Madabhushi, A., Shah, P., Spitzer, M., and Zhao, S. (2019). Applications of machine learning in drug discovery and development. Nature Reviews Drug Discovery.

Vaswani, A., Shazeer, N., Parmar, N., Uszkoreit, J., Jones, L., Gomez, A. N., Kaiser, Ł., and Polosukhin, I. (2017). Attention is all you need. Advances in neural information processing systems, 30.

Vignac, C., Krawczuk, I., Siraudin, A., Wang, B., Cevher, V., and Frossard, P. (2022). Digress: Discrete denoising diffusion for graph generation.

Weininger, D. (1990). Smiles. 3. depict. graphical depiction of chemical structures. Journal of chemical information and computer sciences, 30(3):237–243.

Wu, X., Zhang, Q., Wu, Y., Wang, H., Li, S., Sun, L., and Li, X. (2021). F3a-gan: Facial flow for face animation with generative adversarial networks. IEEE Transactions on Image Processing, 30:8658–8670.

Xu, M., Yu, L., Song, Y., Shi, C., Ermon, S., and Tang, J. (2022). Geodiff: A geometric diffusion model for molecular conformation generation.

Zang, C. and Wang, F. (2020). Moflow: An invertible flow model for generating molecular graphs. pages 617–626. ACM.

