## Supplementary Material for "TumFlow: An AI Model for Predicting New Anticancer Molecules"

*Preprint*, 2024, 1–7
doi: NA

Supplementary Material

### TumFlow: An AI Model for Predicting New Anticancer Molecules

**Davide Rigoni,[1,*] Sachithra Yaddehige,[2] Nicoletta Bianchi,[3] Alessandro Sperduti,[4] Stefano Moro[1] and Cristian Taccioli[2]**

[1]Molecular Modelling Section (MMS), Department of Pharmaceutical and Pharmacological Sciences, University of Padova, Via Marzolo 5, 35131 , Padova, Italy, [2]Department of Animal Medicine, Production and Health, University of Padova, Viale dell'Università 16, 35020, Legnaro, Italy, [3]Department of Translational Medicine, University of Ferrara, Via Luigi Borsari 46, 44121, Ferrara, Italy and [4]Department of Mathematics "Tullio Levi-Civita", University of Padova, Via Trieste 63, 35131, Padova, Italy

##### Abstract

This document contains the Supplementary Material of the article *TumFlow: An AI Model for Predicting New Anticancer Molecules* submitted to the *Bioinformatics, 2024.*

**Key words:** Generative Model, Anticancer Molecules, Melanoma, SK-MEL-28

#### S1. Trends in Drug Development

The drug discovery and design process are complex and resource intensive, often extending over 10-20 years with costs exceeding $2 billion (Zang and Wang, 2020; Harrer et al., 2019). Figure S1 presents the number of FDA approved drugs per year (Mullard, 2022), highlighting the small increase in approvals, despite every year increasing investments in research and development (Statista, 2023) visible in Figure S2.

#### S2. Preliminary Data Analysis

Figure S3 presents a preliminary data analysis performed on the NCI-60 dataset in the context of SK-MEL-28 melanoma cells. Specifically, it reports a violinplot for the GI50, LC50, IC50, and TGI scores highlighting the values of four clinical drugs: (i) Dacarbazine, (ii) Vemurafenib, (iii) Trametinib, and (iv) Cobimetnib. The figure reveals that molecules used clinically *show better representation* according to the GI50 scores. Clearly, the GI50 scores offer a more accurate representation of clinical drugs since they exhibit a more evenly distributed pattern and better distinguish the effects of various drugs.

#### S3. Background: Normalizing Flow Framework

The normalizing flow framework is based on the simple idea of transforming complex data $\boldsymbol{x} \sim \mathbb{P}_x(\boldsymbol{X})$ to simpler representation $\boldsymbol{z} \sim \mathbb{P}_z(\boldsymbol{Z})$ through a series of $l$ invertible and differentiable functions $f = f_l \circ f_{l-1} \circ ... \circ f_1$ as $\boldsymbol{x} = f(\boldsymbol{z})$. The multivariate probability $\mathbb{P}_x(\boldsymbol{X})$ represents the unknown probability distribution that has generated the data $\boldsymbol{x}$ in the dataset $\mathcal{D}$, while the multivariate probability distribution $\mathbb{P}_z(\boldsymbol{Z})$ is chosen to be manageable and often is a standard gaussian normal distribution $\mathcal{N}(0, \boldsymbol{I})$, with zero mean and identity matrix as covariance matrix. Through these invertible transformations, it is possible to transit from a set of original high dimensional data (molecules) to a more manageable and simpler latent space, and vice versa. Think of this process as reordering: starting from a chaotic box full of various objects, the model reorganizes them into a more orderly and comprehensible box (the latent space).

In mathematical terms, using the *change of variable formula*, the transformation can be expressed as:

$$\mathbb{P}_x(\boldsymbol{X}) = \mathbb{P}_z(\boldsymbol{Z}) \cdot \left| \det \frac{\partial \boldsymbol{Z}}{\partial \boldsymbol{X}} \right|,$$

where $\mathbb{P}_x(\boldsymbol{X})$ is the probability of the original data distribution and $\mathbb{P}_z(\boldsymbol{Z})$ that of the latent space. The $l$ functions are learned through the calculation of log-likelihood, a measure indicating how accurately the functions represent the data in the training set:

$$\log \mathbb{P}_x(\boldsymbol{X}) = \log \mathbb{P}_z(\boldsymbol{Z}) + \log \left| \det \frac{\partial \boldsymbol{Z}}{\partial \boldsymbol{X}} \right|;$$

$$= \log \mathbb{P}_z\left(f^{-1}(\boldsymbol{X})\right) + \log \left| \det \frac{\partial f^{-1}(\boldsymbol{X})}{\partial \boldsymbol{X}} \right|.$$

To generate a new data point $\hat{\boldsymbol{x}} \sim \mathbb{P}_x(\boldsymbol{X})$, when the function $f$ is learned, it is sufficient to sample a new point $\hat{\boldsymbol{z}}$ from the

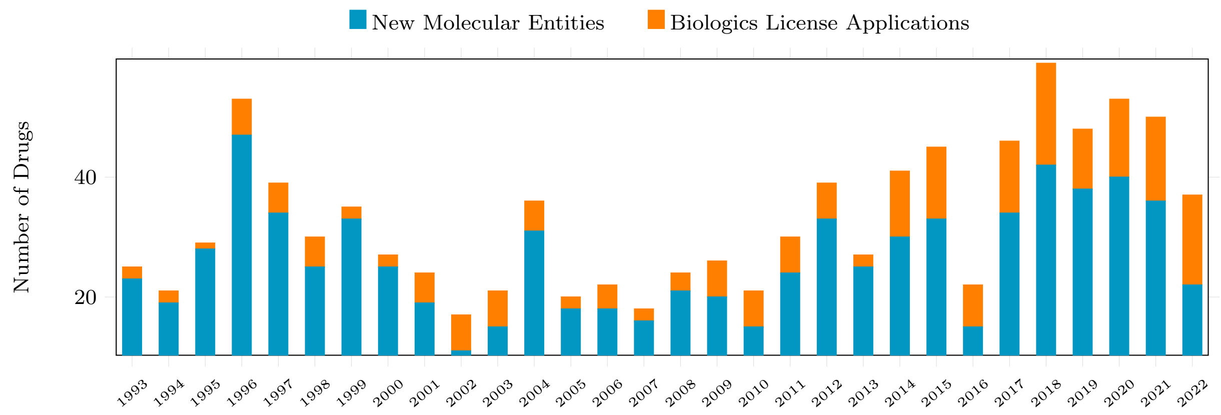


Fig. S1: Number of FDA-approved drugs per year. Products like vaccines and gene therapies are not included in this drug count. The plot depicts a trend where the number of approved molecules shows only a marginal increase over the years.

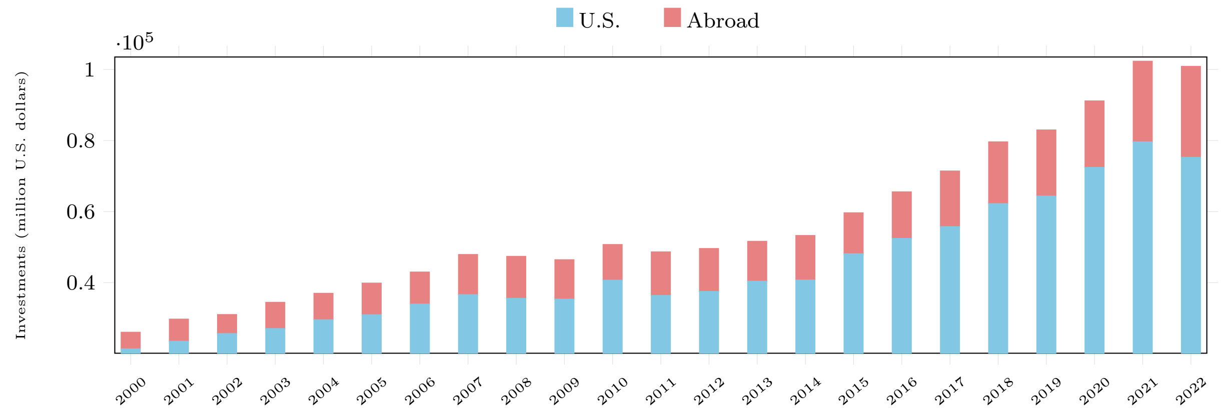


Fig. S2: Spending of the U.S. pharmaceutical industry, by PhRMA member companies, on research and development at home and abroad. Over the years, there has been a substantial increase in financial investments dedicated to research and development.

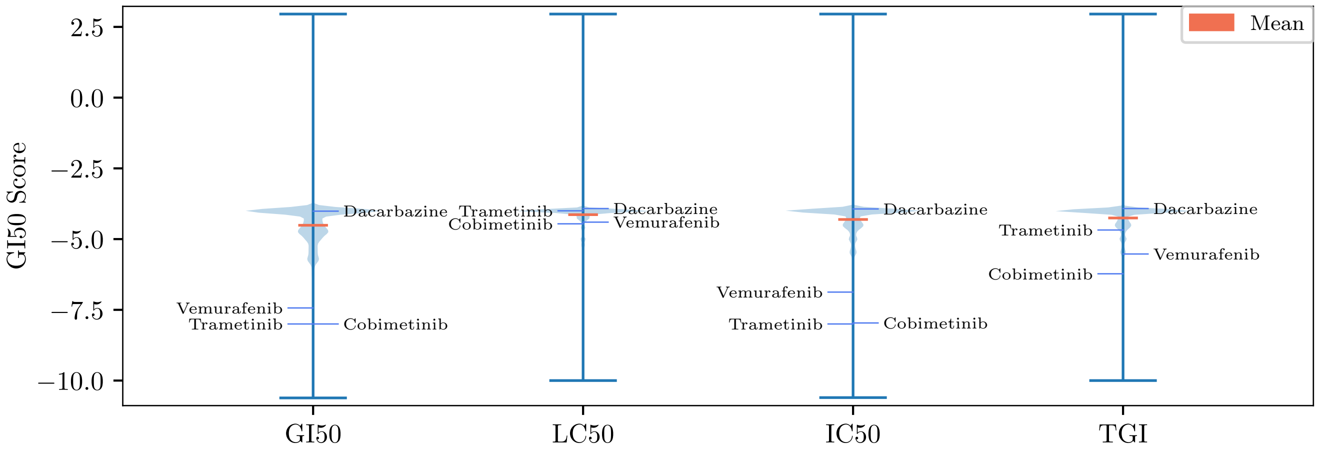


Fig. S3: Violinplot of the GI50, LC50, IC50, and TGI scores available from the National Cancer Institute database highlighting the values of four clinical drugs: (i) Dacarbazine, (ii) Vemurafenib, (iii) Trametinib, and (iv) Cobimetnib. The mean is reported with the orange colour. The GI50 scores offer a more accurate representation of clinical drugs since they exhibit a more evenly distributed pattern and better distinguish the effects of various drugs.

probability distribution $\mathbb{P}_z(\boldsymbol{Z})$ and then to obtain $\hat{\boldsymbol{x}}$ as:

$$\hat{\boldsymbol{x}} = f(\hat{z}).$$

A crucial component of normalizing flow models is the invertible coupling layers that aim to implement the invertible differentiable function $f = f_l \circ f_{l-1} \circ ... \circ f_1$. These layers allow a wide range of transformations, making the model flexible and reliable, yet allowing the exploitation of the backpropagation algorithm essential to train neural networks. Specifically, they are required to be expressive enough to transform the data and allow efficient computation of the Jacobian determinant of the matrix.

The literature presents several formulations of coupling layers, such as additive coupling layers (Dinh et al., 2014) and affine coupling layers (Dinh et al., 2016; Kingma and Dhariwal, 2018). The core idea of these approaches is to divide input data into two groups, with one group conditioning the transformation applied to the second group. Mathematically, let $f_i^{-1} \forall i \in \{1, ..., l\}$ composing $f^{-1}$ be an affine coupling layer defining a function $f_i^{-1} : \mathbb{R}^d \to \mathbb{R}^d$, where $\boldsymbol{z}^{i-1} = f_i^{-1}(\boldsymbol{z}^i)$, $\boldsymbol{z} = \boldsymbol{z}^0$, $\boldsymbol{x} = \boldsymbol{z}^{l+1}$, and $d =| \boldsymbol{z}^i |$ is the length of vector $\boldsymbol{z}^i$. Then, the dimensions of the vector in the input are split into two groups by a chosen dimension $\widetilde{d}$, such that:

$$\boldsymbol{z}^{i-1} = \begin{cases} \boldsymbol{z}^{i-1}_{[1:\widetilde{d}]} & = \boldsymbol{z}^{i}_{[1:\widetilde{d}]}; \\ \boldsymbol{z}^{i-1}_{[\widetilde{d}+1:d]} & = \boldsymbol{z}^{i}_{[\widetilde{d}+1:d]} \odot exp\left(s\left(\boldsymbol{z}^{i}_{[1:\widetilde{d}]}\right)\right) + t\left(\boldsymbol{z}^{i}_{[1:\widetilde{d}]}\right); \end{cases}$$

where $\odot$ is the element-wise product, $s : \mathbb{R}^{\widetilde{d}} \to \mathbb{R}^{d-\widetilde{d}}$ is the scale function, and $t : \mathbb{R}^{\widetilde{d}} \to \mathbb{R}^{d-\widetilde{d}}$ is the translation function. Note that arbitrary complex neural networks can be employed to implement both functions $s$ and $t$ and that the application of several affine coupling layers in sequence, which alternate the data partitions to transform, ensures that every dimension undergoes a transformation at some point during the process.

#### S4. TumFlow: Additional Details

This section presents more details on the TumFlow model. Specifically, the sanitization of the molecules, the model selection and the implementation details will be clarified in the following.

##### S4.1. Sanitization of the Generated Molecules

The molecule graphs predicted by *TumFlow* are not free from errors, and without any precaution, the direct assembly of the molecule atoms and bonds from the generated graph may lead to chemically invalid molecules. For this reason, *TumFlow* employs a post-processing algorithm aiming, whenever possible, to satisfy the chemical constraints derived by atom valencies. More precisely, this process is defined for an atom $i \in G$ as:

$$\sum_{j,e} \boldsymbol{E}_{i,j,e} \cdot f_e(e) \leq f_{av}(i) + f_{fc}(i),$$

where $f_e(e)$ is the function that returns the number of electrons of atom $i$ constrained by a bond type $e$, while $f_{av}(i)$ and $f_{fc}(i)$ are the functions that return the valence and formal charge of atom $i$, respectively. Only for atom types N and S, the formal charge is allowed to be 1, otherwise, it is 0. This allows the generation in the output of molecules with atom $N^+$ and $S^+$. Upon compliance with this rule, RDKit[1] is used to assemble the molecule.

[1] https://www.rdkit.org/

##### S4.2. Model Selection

*TumFlow* is trained on the training set, while the model selection is done by looking at the loss values in the validation set. Precisely, the best model is the one that achieved the lower negative log-likelihood loss in the validation during the training of the normalizing flow model and the lower mean squared error loss in the validation during the training of the optimizer.

The search for a performing *TumFlow* model started by using the values of the hyper-parameters defined by MoFlow. Subsequently, based on the observed loss on the validation set, these values were iteratively adjusted by varying the number and dimensions of the layers within the neural network. When adding depth to the networks did not improve the validation loss anymore, a grid search to find the best gradient descend steps to adopt during the generation of new molecules was considered with the following values: $\{0.001, 0.0001, 0.0005\}$. The final batch size is selected among $\{64, 128, 256, 512\}$.

In the exploration of optimal hyperparameters, diverse values were considered for the number of coupling layers in both $f_{\mathcal{E}}$ and $f_{\mathcal{V}|\mathcal{E}}$. Specifically, the examined options for the number of layers $l_{\mathcal{E}}$ were $\{7, 10, 20\}$, while for $l_{\mathcal{V}|\mathcal{E}}$, they were $\{38, 100, 130\}$. The functions $s^i_{\mathcal{E}}$ and $t^i_{\mathcal{E}}$ were implemented using a sequence of two $3 \times 3$ 2D convolutional networks, with dimensions chosen from $\{128, 256, 512\}$. For the graph neural networks employed in $s^i_{\mathcal{V}|\mathcal{E}}$ and $t^i_{\mathcal{V}|\mathcal{E}}$, the feature dimensions were selected from $\{128, 256\}$. Additionally, the functions $s^i_{\mathcal{V}|\mathcal{E}}$ and $t^i_{\mathcal{V}|\mathcal{E}}$ were implemented as MLPs with two hidden layers with dimensions in $\{128, 256, 512\}$.

Due to limitations in the available computational resources, the fine-tuning of the other parameters was not performed.

##### S4.3. Implementation Details

The experiments were performed on a cluster with 64GB of RAM and an A100 GPU with 40GB of memory. The training set is formed by $41,556$ molecules, while the validation set is formed by $5,000$ molecules. Both training and validation molecules are composed of a maximum of 80 atoms. The random noise selected to transform discrete tensors to continue tensors is drawn from the uniform distribution $U[0, 0.6]$. The best normalizing flow model is obtained at epoch 3, using a batch size of 256 examples and a learning rate of 0.001. In the end, the best model was achieved with $l_{\mathcal{E}} = 7$ and $l_{\mathcal{V}|\mathcal{E}} = 130$ coupling layers. Concerning $f_{\mathcal{E}}$, all the convolutional neural networks are configured with 128 neurons. Regarding the graph neural networks and the MLPs implemented in $f_{\mathcal{V}|\mathcal{E}}$, the chosen hidden dimension is 128, which has demonstrated the best performance.

After the successful training of the main normalizing flow architecture, the optimization network is subsequently trained. During this final training phase, the weights of the flow architecture are frozen, i.e., they are not finetuned. The best optimizer was obtained at epoch 8, using a batch size of 256 examples, a learning rate of 0.001 and considering unnormalized GI50 scores. In the generation of new molecules, 100 optimization steps were considered for each molecule, with two different gradient descent steps: $\{0.001, 0.005\}$.

During the implementation of the affine coupling layers $\Phi_i$, the actnorm2D technique is utilized to normalize the channel features across a batch of atom matrices. Similarly, the one-dimensional version is applied in the implementation of the coupling layers $\Psi_i$. Furthermore, to increase the number of channels to split

**Table S1.** List of antitumoral drugs considered in this work, which are known in clinical treatments against melanoma. The coloured lines highlight the molecules used as a starting point to obtain the results presented in Section 3.2 of the main article.

| NSC ID | Name | Canonical SMILES |
|---|---|---|
| 8806 | Alkeran | C1=CC(=CC=C1CC(C(=O)O)N)N(CCCl)CCCl |
| 768068 | Cobimetinib | C1CCNC(C1)C2(CN(C2)C(=O)C3=C(C(=C(C=C3)F)F)NC4=C(C=C(C=C4)I)F)O |
| 764134 | Dabrafenib Mesylate | CC(C)(C)C1=NC(=C(S1)C2=NC(=NC=C2)N)C3=C(C(=CC=C3)NS(=O)(=O)C4=C(C=CC=C4F)F)F |
| - | Encorafenib | CC(C)N1C=C(C(=N1)C2=C(C(=CC(=C2)Cl)NS(=O)(=O)C)F)C3=NC(=NC=C3)NCC(C)NC(=O)OC |
| 758246 | Trametinib | CC1=C2C(=C(N(C1=O)C)NC3=C(C=C(C=C3)I)F)C(=O)N(C(=O)N2C4=CC(=CC=C4)NC(=O)C)C5CC5 |
| 720033 | Interferon alfa-2B | CCCN(CCOC1=C(C=C(C=C1Cl)Cl)Cl)C(=O)N2C=CN=C2 |
| 761431 | Vemurafenib | CCCS(=O)(=O)NC1=C(C(=C(C=C1)F)C(=O)C2=CNC3=NC=C(C=C23)C4=CC=C(C=C4)Cl)F |
| 45388 | Dacarbazine | CN(C)NN=C1C(=NC=N1)C(=O)N |
| - | Binimetinib | CN1C=NC2=C1C=C(C(=C2F)NC3=C(C=C(C=C3)Br)F)C(=O)NOCCO |

and transform, each coupling layer also applies a reshaping transformation to the tensors to obtain a different view of it.

More details about the *TumFlow* code, along with the datasets used and produced, are available in an online public repository[2].

#### S5. Clinical Atitumoral Drugs

Table S1 reports the list of antitumoral drugs considered in this work, which are known in clinical treatments against melanoma. Each drug is reported with its "NSC ID" code, name and canonical SMILES. The coloured lines highlight the molecules used as a starting point to obtain the results presented in Section 3.2 of the main article.

#### S6. Generated Molecules

This section presents the molecules generated with *TumFlow*. In particular, Table S2 and Table S3 report the lists of novel molecules visible in Figure 3 and Figure 4 of the main manuscript, respectively. The molecules reported in these tables are generated from those in the NCI-60 dataset. On the contrary, Table S4 displays the list of molecules visible in Figure 5 of the main article, that are generated from those used in clinic for the treatment of melanoma. The coloured lines highlight the molecules already present on PubChem[3].

#### S7. TumFlow Limitations

The use of GI50 data, extracted from the NCI-60 project, utilizing the TumFlow model has yielded significant findings. The results demonstrated that TumFlow, with its advanced architecture and capacity to handle complex discrete data, could discern key patterns and correlations between molecular structures and their antitumoral effectiveness. This indicates that the model can be employed not only for predicting the GI50 efficacy score but also for designing new compounds with potential therapeutic properties. However, it is crucial to note that, despite promising results, the model requires further validation with additional experimental data. Moreover, the complexity of molecular interactions and biological mechanisms implies that computational findings must be approached with caution and corroborated through comprehensive experimental studies.

In this regard, *TumFlow* has shown a propensity to produce energetically unstable structures and highly complex molecules, presenting synthesis challenges. This is not surprising given the difficulty of predicting novel energetically stable molecules that can be synthesised with modern technology. Figure S4 illustrates some examples of generated molecules that, while adhering to valence laws, show certain issues regarding the *TumFlow* generated structures. Specifically, the initial seven molecules exhibit energetically unstable structures primarily due to a group of atoms that cannot be synthesized. Examples eight to twelve represent instances where *TumFlow* failed to accurately predict aromaticity. The remaining structures predominantly consist of macrocycles, amplifying the challenges in the synthesis process. However, despite the difficult synthesis, *TumFlow* associates them with the predicted GI50 score, suggesting that these molecules exhibit a high predicted antitumoral efficacy.

[2] https://github.com/drigoni/TumFlow

[3] https://pubchem.ncbi.nlm.nih.gov/

**Table S2.** List of novel molecules generated from those in the NCI-60 dataset. The molecule structures of the following SMILES strings are visible in Figure 3 of the main manuscript. The coloured lines highlight the molecules already present on PubChem.

| Starting Molecule | Generated Molecule |
|---|---|
| CC(C)C12OC1C1OC13C1(C)CCC4=C(COC4=O)C1CC1OC13C2O | COC1C2(C(C)C)OC2C2OC23C2(C)CCC4=C(COC4=O)C2CC2OC213 |
| CC(C)C12OC1C1OC13C1(C)CCC4=C(COC4=O)C1CC1OC13C2O | CCOC1C2(C(C)C)OC2C2OC23C2(C)CCC4=C(COC4=O)C2CC2OC213 |
| CC(C)C12OC1C1OC13C1(C)CCC4=C(COC4=O)C1CC1OC13C2O | CCCOC1C2(C(C)C)OC2C2OC23C2(C)CCC4=C(COC4=O)C2CC2OC213 |
| CC(C)C12OC1C1OC13C1(C)CCC4=C(COC4=O)C1CC1OC13C2O | CCCCCOC1C2(C(C)C)OC2C2OC23C12OC2CC12OC4OCC1=C4CCC23C |
| COc1cc2c(c(OC)c1O)-c1ccc(SC)c(=O)cc1C(NC(C)=O)CC2 | COc1cc2c(c(OC)c1O)-c1ccc(SC)c(=O)cc1C(NC1CO1)CC2 |
| COc1cc2c(c(OC)c1O)-c1ccc(SC)c(=O)cc1C(NC(C)=O)CC2 | COc1cc2c(c(OC)c1O)-c1ccc(SC)c(=O)cc1C1NC(C)OC1C2 |
| COc1cc2c(c(OC)c1O)-c1ccc(SC)c(=O)cc1C(NC(C)=O)CC2 | COc1cc2c(c(OC)c1O)-c1ccc(SC)c(=O)cc1C1NC(C)OC1C2C |
| COc1cc2c(c(OC)c1O)-c1ccc(SC)c(=O)cc1C(NC(C)=O)CC2 | CCCC1c2cc(OC)c(O)c(OC)c2-c2ccc(SC)c(=O)cc2C2NC(C)OC12 |

**Table S3.** List of novel molecules generated from those in the NCI-60 dataset. The molecule structures of the following SMILES strings are visible in Figure 4 of the main manuscript.

| Starting Molecule | Generated Molecule |
|---|---|
| C=Cc1cc(OC)c2c(c1)c(=O)oc1c2cc(OC)c2c(O)ccc(C3OC(C(C)O)C(O)C3O)c21 | C=Cc1cc(OC)c2c(c1)c(=O)oc1c2cc(OC)c2c(O)ccc(C3OC(C4CO4)C(O)C3O)c21 |
| CCOc1ccc(C=Cc2cc(OC)c(OC)c(OC)c2)cc1 | CCOc1ccc2ccc3cc(OC)c(OC)c(OC)c3c2c1 |
| Cc1ccnc2[nH]c(-c3cccc4ccccc34)cc(=O)c12 | Cc1cnc2[nH]c(-c3cccc4ccccc34)cc(=O)c2c1C |
| Cc1ccnc2[nH]c(-c3cccc4ccccc34)cc(=O)c12 | CCCCCCC1C(C)=CN=C2NC(c3cccc4ccccc34)=CC(=O)C21 |
| COc1cc2c(c(OC)c1OC)-c1ccc(SC)c(=O)cc1C(NCc1ccc([N+](=O)[O-])cc1)CC2 | COc1cc2c(c(OC)c1OC)-c1ccc(SC)c(=O)cc1C(NCc1ccc([N+](=O)O)cc1)CC2 |
| COC(=O)CCNc1c2ccccc2nc2cccc([N+](=O)[O-])c12 | COC(=O)CCNc1c2ccccc2nc2cccc([N+](=O)O)c12 |
| CCOC(=O)Nc1cc2c(c(N)n1)N=C(c1ccc(OC)cc1)CN2 | CCOC(=O)Nc1cc2c(c(N)n1)N=C(c1ccc(OC)cc1)CN2C |
| CCOC(=O)Nc1cc2c(c(N)n1)N=C(c1ccc(OC)cc1)CN2 | CCOC(=O)Nc1cc2c(c(N)n1)N=C(c1ccc(OC)cc1)CN2CC |
| C=C1COC(O)(C2OCOC(C3CC(=O)NC(=O)C3)C2O)C1C | C=C1COC2(O)C3OCOC(C4CC(=O)NC(=O)C4)C3OC12C |
| C=C1COC(O)(C2OCOC(C3CC(=O)NC(=O)C3)C2O)C1C | C=C1COC2(O)C3OCOC(C4CC(=O)NC(=O)C4)C3OC12CCCCC |

**Table S4.** List of novel molecules generated from those used in clinic for the treatment of melanoma. The molecule structures of the following SMILES strings are visible in Figure 5 of the main manuscript. The coloured lines highlight the molecules already present on PubChem.

| Drug Name | Starting Molecule | Generated Molecule |
|---|---|---|
| Alkeran | C1=CC(=CC=C1CC(C(=O)O)N)N(CCCl)CCCl | COC(=O)C(N)Cc1ccc(N(CCCl)CCCl)cc1 |
| Alkeran | C1=CC(=CC=C1CC(C(=O)O)N)N(CCCl)CCCl | NC1Cc2ccc(N(CCCl)CCCl)cc2C1O |
| Alkeran | C1=CC(=CC=C1CC(C(=O)O)N)N(CCCl)CCCl | COC1(O)c2cc(N(CCCl)CCCl)ccc2CC1N |
| Alkeran | C1=CC(=CC=C1CC(C(=O)O)N)N(CCCl)CCCl | C=C(CCCCCCCCC)C1(O)c2cc(N(CCCl)CCCl)ccc2CC1N |
| Interferon alfa-2B | CCCN(CCOC1=C(C=C(C=C1Cl)Cl)Cl)C(=O)N2C=CN=C2 | CCCN(CCOc1c(Cl)cc(Cl)cc1Cl)C(=O)N1C=CNC1 |
| Interferon alfa-2B | CCCN(CCOC1=C(C=C(C=C1Cl)Cl)Cl)C(=O)N2C=CN=C2 | CCCC1NC=CN1C(=O)N(CCC)CCOc1c(Cl)cc(Cl)cc1Cl |
| Interferon alfa-2B | CCCN(CCOC1=C(C=C(C=C1Cl)Cl)Cl)C(=O)N2C=CN=C2 | CCCCCC1NC=CN1C(=O)N(CCC)CCOc1c(Cl)cc(Cl)cc1Cl |
| Interferon alfa-2B | CCCN(CCOC1=C(C=C(C=C1Cl)Cl)Cl)C(=O)N2C=CN=C2 | CCCCCCC1NC=CN1C(=O)N(CCC)CCOc1c(Cl)cc(Cl)cc1Cl |

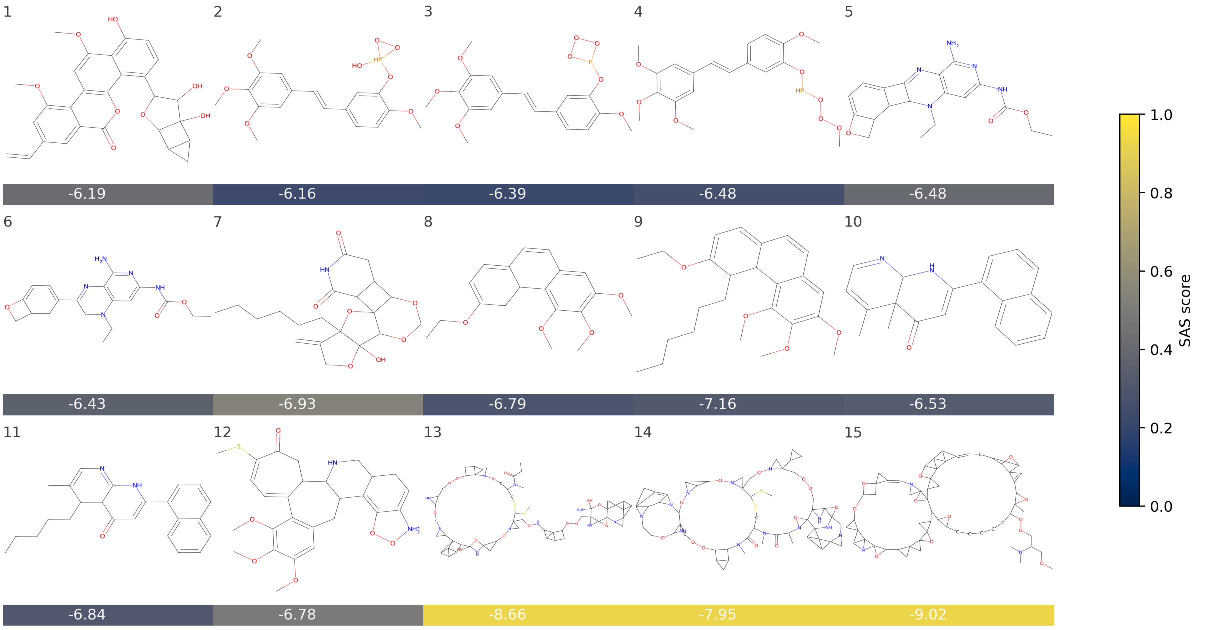


Fig. S4: This grid underscores a limitation of *TumFlow*, i.e., its tendency to generate energetically unstable structures or molecules that are challenging to synthesize. The score reported under each generated molecule represents the *TumFlow* predicted GI50 score, while the colour conveys the normalized SAS score. The value on the molecule top-left box reports the molecule index.

#### References

Dinh, L., Krueger, D., Bengio, Y., and Courville, A. (2014). Nice: Non-linear independent components estimation. *Advances in Neural Information Processing Systems*, pages 2224–2232.

Dinh, L., Sohl-Dickstein, J., and Bengio, S. (2016). Density estimation using real nvp. In *International Conference on Learning Representations*.

Harrer, S., Shah, P., Antony, B., and Hu, J. (2019). Artificial intelligence for clinical trial design. *Trends in pharmacological sciences*, 40:577–591.

Kingma, D. P. and Dhariwal, P. (2018). Glow: Generative flow with invertible 1x1 convolutions. *Advances in neural information processing systems*, 31.

Mullard, A. (2022). 2021 fda approvals. *Nature reviews. Drug discovery*.

Statista (2023). U.S. pharmaceutical industry spending on research and development 1990-2022. https://www.statista.com/statistics/265090/us-pharmaceutical-industry-spending-on-research-and-development/.

Zang, C. and Wang, F. (2020). Moflow: An invertible flow model for generating molecular graphs. pages 617–626. ACM.
